# The Dimer-dependent Catalytic Activity of RAF Family Kinases Is Revealed Through Characterizing Their Oncogenic Mutants

**DOI:** 10.1101/338293

**Authors:** Jimin Yuan, Wan Hwa Ng, Paula Y.P. Lam, Yu Wang, Hongping Xia, Jiajun Yap, Shou Ping Guan, Ann S.G. Lee, Mei Wang, Manuela Baccarini, Jiancheng Hu

**Affiliations:** Division of Cellular and Molecular Research; Division of Medical Sciences, National Cancer Centre Singapore; 11 Hospital Drive, 169610, Singapore.; Cancer and Stem Cell Program; Office of Clinical & Academic Faculty Affairs, Duke-NUS Medical School; 8 College Road, 169857, Singapore.; Department of Physiology, National University of Singapore; 2 Medical Drive. 117597, Singapore; Max F. Perutz Laboratories, University of Vienna, Doktor-Bohr-Gasse 9, 1030 Vienna, Austria.

**Keywords:** protein kinase, signal transduction, oncogenic kinase mutations, drug resistance, cancer

## Abstract

Although extensively studied for three decades, the molecular mechanisms that regulate the RAF/MEK/ERK kinase cascade remain ambiguous. Recent studies identified the dimerization of RAF as a key event in the activation of this cascade. Here, we show that in-frame deletions in the β3-αC loop activate ARAF as well as BRAF and other oncogenic kinases by enforcing homodimerization. By characterizing these RAF mutants, we find that ARAF has less allosteric and catalytic activity than the other two RAF isoforms, which arises from its non-canonical APE motif. Further, these RAF mutants exhibit a strong oncogenic potential, and a differential inhibitor resistance that correlates with their dimer affinity. Using these unique mutants, we demonstrate that active RAFs, including the BRAF(V600E) mutant, phosphorylate MEK in a dimer-dependent manner. This study characterizes a special category of oncogenic kinase mutations, and elucidates the molecular basis that underlies the differential ability of RAF isoforms to stimulate MEK-ERK pathway. Further, this study reveals a unique catalytic feature of RAF family kinases that can be exploited to control their activities for cancer therapies.

## Introduction

The Ras/RAF/MEK/ERK signaling plays a crucial role in cell proliferation, survival, and differentiation ^1, 2^. Aberrant activation of this kinase cascade causes developmental disorders and cancers ^3-5^. Genetic alterations that hyperactivate the RAF/MEK/ERK kinase cascade exist in >40% of cancers. To target this kinase cascade for cancer therapy, both RAF inhibitors and MEK inhibitors have been developed and applied to clinical treatment ^6-8^. Unfortunately, their efficacy is limited by either intrinsic or rapidly acquired resistance. Understanding the regulation of the RAF/MEK/ERK kinase cascade could help us to design strategies to circumvent this resistance and develop more effective inhibitors for cancer treatment.

The RAF kinases CRAF, BRAF and ARAF are a core component of the RAF/MEK/ERK kinase cascade. Dimerization among RAF isoforms is a key event in triggering the RAF/MEK/ERK kinase cascade ^9-18^, which not only turns on the kinase activity of RAF but also facilitates the activation of MEK by RAF ^19^. Mechanistic studies have shown that the two protomers play distinct roles in RAF dimers: one functions as an allosteric activator to facilitate the assembly of an active conformation in the other, which acts as a receiver to catalyze the phosphorylation of substrates ^20^. Distinct molecular traits between BRAF and CRAF results in their differential ability to turn on the RAF/MEK/ERK kinase cascade by dimerization-driven transactivation ^21^. Whether ARAF can be activated by dimerization and its role in this process are unclear.

The dimerization of RAF kinase occurs in both physiological and pathological conditions, which can be induced by active Ras ^13, 17^, inhibitor binding ^9, 11, 12, 14^, gene fusions ^22-25^, and alternative splicings ^26^. Active Ras-induced homo/hetero-dimerization of RAF kinase is responsible not only for the pathway activation by physiological agonists but also for its hyperactivation by genetic alterations upstream of RAF in carcinogenesis ^13, 17, 27^. Active Ras-induced RAF dimerization can be further enhanced by RAF inhibitors, which accounts for the paradoxical effect of RAF inhibitors in cancer therapy ^11, 12, 14^. It has been speculated that the RAF kinases have a close conformation in which the N-terminus docks on the carboxyl-terminal kinase domain ^28^. Active Ras binds to the N-terminus of RAF kinases, which breaks their close conformation and thus facilitates their dimerization via kinase domain. On the other hand, RAF inhibitors could alter the conformation of RAF kinase domain once loaded, which not only creates a dimer-favored conformation but also relives the inhibitory interaction between N-terminus and kinase domain ^29^. The inhibitory effect of N-terminus on RAF dimerization could also be abolished by gene fusions or alternative splicing of mRNA. Chromosome translocations that lead to fusions of variable genes to the kinase domain of BRAF or CRAF have been extensively reported in cancers ^22-25^, while the alternative splicings that partially delete the N-terminus of BRAF(V600E) and thus enhance the dimerization of BRAF(V600E) isoforms have been shown as one of important mechanisms that underlie RAF inhibitor resistance in cancer therapy ^26^. Recently, some RAF mutants (ARAF and BRAF) with in-frame deletions in the β3-αC loop have been reported as potential oncogenic drivers ^30-34^, although whether they are activated through enhanced dimerization remains unknown ^30, 31, 33^ or controversial ^32,34^.

In this study, we characterized the RAF mutants with in-frame deletions in the β3-αC loop, and found that both ARAF and BRAF mutants were activated by enhanced dimerization. Further, we showed that the limited allosteric and catalytic activities of ARAF arose from its non-canonical APE motif that leads to a lower propensity of dimerization in contrast to BRAF and CRAF. Finally, we used active RAF mutants with different dimerization properties as an efficient tool to investigate whether the dimerization of RAF after activation is required for its catalytic activity and demonstrated that active RAFs including BRAF(V600E) phosphorylated MEK in a dimer-dependent manner. Our data clarifies how in-frame β3-αC loop deletions trigger RAF family kinases, reveals the molecular basis underlying the differential ability of RAF isoforms to stimulate MEK-ERK signaling, and illustrates a key step in the activation of the RAF/MEK/ERK kinase cascade.

## Results

### Deletion of Q347A348 activates ARAF by enforcing dimerization

By virtue of its apparent low activity and rare mutations in cancer genomes, the molecular mechanism regulating ARAF and its role in oncogenesis are ill-defined. Recently, Nelson et al. identified an active ARAF mutant with Q347A348 deletion and F351L conversion in patients with Langerhans cell histiocytosis ^31^. To confirm this finding and to decipher the molecular basis of ARAF activation by this compound mutation, we expressed wild-type ARAF, and its Q347A348del mutants (ΔQA) with or without F351L mutation in 293T cell, and found that Q347A348del mutant was able to stimulate the MEK-ERK pathway independent of F351L status (Figure 1A), suggesting a dominant role of the Q347A348 deletion in the activation of ARAF. Similar to other constitutively-active mutants of RAF kinases, a co-expression of a dominant-negative RAS mutant (N17RAS) with ARAF(ΔQA) did not affect its activity in 293T cells (Figure 1B), indicating that ARAF(ΔQA) is a constitutively-active mutant and does not require upstream stimuli to trigger its catalytic activity. This notion was further validated by the finding that a stable expression of ARAF(ΔQA) in wild-type, BRAF-/-, and CRAF-/- fibroblasts activated the MEK-ERK pathway and transformed cells independent of endogenous RAF molecules(Figure S1A and 1C). Moreover, the shRNA-mediated knockdown of either CRAF in BRAF-/- fibroblasts or BRAF in CRAF-/- fibroblasts had no effect on the ability of ARAF(ΔQA) to stimulate downstream signaling (Figure S1B).

**Figure 1.**
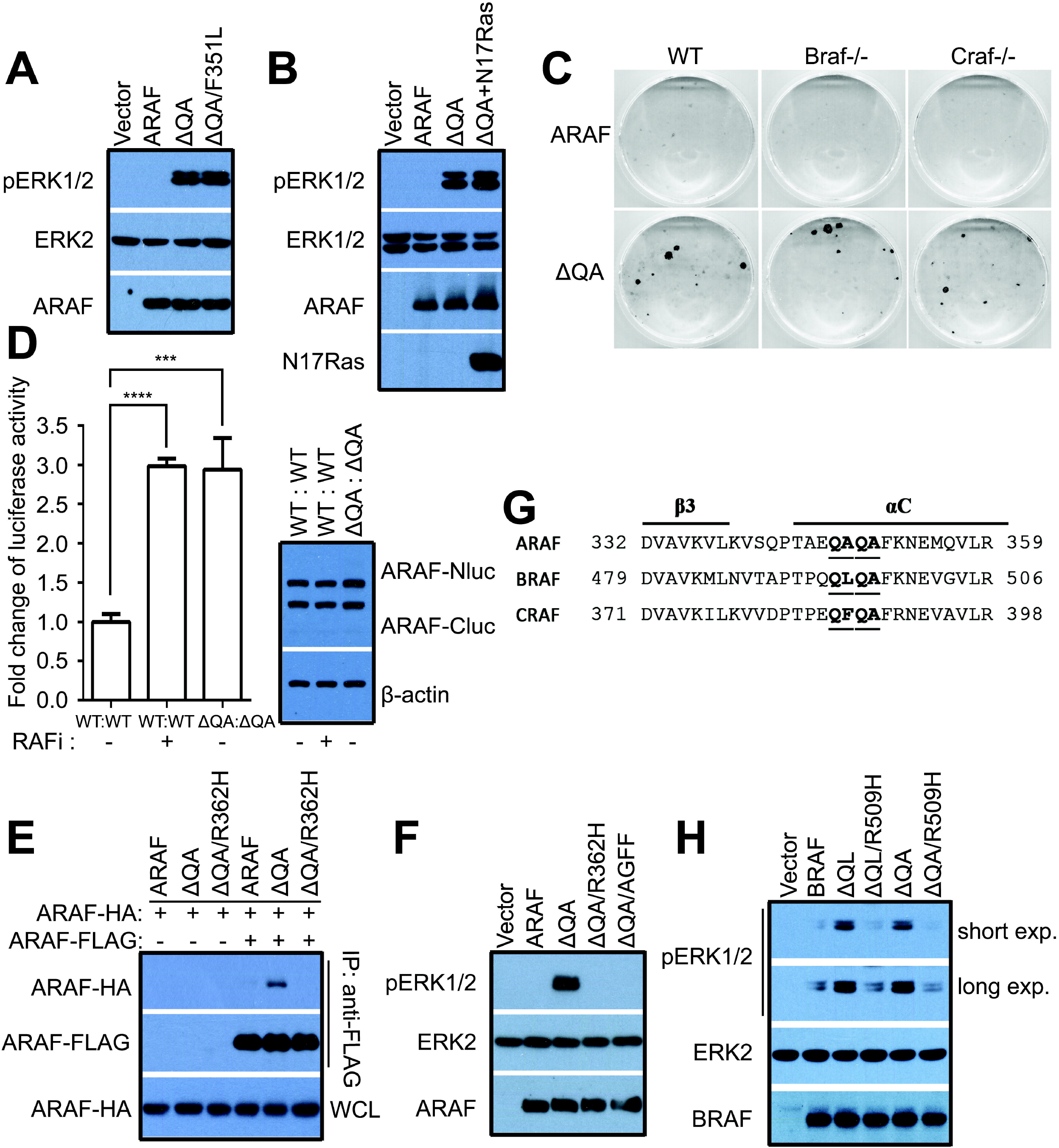
The Q347A348 deletion activates ARAF by enhancing homodimerization. (A) ARAF(ΔQA) and ARAF(ΔQA/ F351L) have equal activity. The activity of ARAF mutants in 293T transfectants was measured by anti-phospho-ERK1/2 immunoblot. (B) The activity of ARAF(ΔQA) does not depend on upstream stimuli. ARAF(ΔQA) was coexpressed with N17Ras in 293T cells, and its activity was measured as in (A). (C) ARAF(ΔQA) has a strong transforming ability independent of endogenous RAF molecules. Foci formation assay of immortalized fibroblasts expressing ARAF(ΔQA) was carried out as described before ^36, 37^. (D-E) ARAF(ΔQA) has an elevated propensity to form homodimers. D, the dimer affinity of ARAF(ΔQA) was measured by using complementary split luciferase assay ^35^. The dimerization of wild-type ARAF induced by 10um Vemurafenib served as a positive control (*n*=5, \*\*\**p*<0.001, \*\*\*\**p*<0.0001). E, the dimerization of ARAF(ΔQA) was evaluated by co-immunoprecipitation assay. ARAF mutants with FLAG- or HA-tag were coexpressed in 293T cells, and immunoprecipitated by anti-FLAG beads and detected by anti-HA immunoblot. To exclude the effect of ERK1/2-mediated feedback on ARAF dimerization, all 293T transfectants in D & E were pre-treated with 20um Tramentinib for 1 hour before measurements. (F) ARAF(ΔQA) is activated by dimerization-driven transactivation. Mutations that disrupt the dimer interface (R362H) or block NtA-phosphorylation (AGFF) abolish the activity of ARAF(ΔQA). The activity of ARAF mutants in 293T transfectants was measured as in (A). (G-H) Homologous deletions activate BRAF in a dimer-dependent manner. G, the sequence alignment of human ARAF, BRAF and CRAF reveals conserved residues in the β3-αC loop. H, the activity of BRAF mutants in 293T transfectants was measured as in (A). All images are representative of at least three independent experiments.

Dimerization of RAF molecules is critical for their activation under variable conditions ^9-17^. We thus investigated whether ΔQA activates ARAF by enhancing dimerization. To do this, we carried out a complementary split luciferase assay ^35^ to measure the dimer affinity of ARAF and its ΔQA mutant. In this assay, the N-terminus and C-terminus of luciferase (hitherto referred to as Nluc and Cluc) were fused to ARAF and co-expressed in cells. RAF dimerization brings the Nluc and Cluc together and reconstitutes the luciferase activity. Thus, the luciferase activity correlates with the amount of dimerized ARAF. As shown in Figure1D, the treatment with RAF inhibitor Vemurafenib, an agonist of RAF dimerization, increased the luciferase activity of 293T cells co-expressing ARAF-Nluc and ARAF-Cluc. In contrast, the 293T cells co-expressing ARAF(ΔQA)-Nluc and ARAF(ΔQA)-Cluc showed a constitutive luciferase activity comparable to that induced by Vemurafenib in wild-type ARAF transfectants, suggesting that ARAF(ΔQA) mutant has an elevated ability to form homodimers. The homodimeriztion of ARAF(ΔQA) in 293T cells can be further verified by co-immunoprecipitation assay. In contrast to its wild-type counterpart, HA-tagged ARAF(ΔQA) could be pull-down by its FLAG-tagged version when co-expressed in 293T cells although the amount is a little (Figure1E). Using the same method, we next evaluated the ability of ARAF(ΔQA) to heterodimerize with BRAF, CRAF and its wild-type counterpart, and found that it barely dimerized with these molecules (Figure S1C).

It has been reported that the dimerization-driven transactivation of RAF molecules might require the interaction of the negatively charged N-terminal acidic motif (NtA motif) with the RKTRH motif in the αC-helix-β4 loop of the other protomer, and mutations that abrogate the negative charge of the NtA motif or disrupt dimer interface block this process ^20^. To determine whether ARAF(ΔQA) is activated through a dimerization-driven transactivation, we mutated its NtA motif (SGYY to AGFF) or its central residue in the dimer interface (R362H), and found that both alterations impaired its activity (Figure 1F), providing additional evidence that deletion of Q347A348 activates ARAF by enhancing homodimerization.

### Homologous deletions of two residues activate BRAF in dimer-dependent manner

Since the Q347A348del activates ARAF by enhancing homodimerization and these two residues are conserved in the β3-αC loop across all RAF isoforms (Figure 1G), we next asked whether a homologous deletion would activate other RAF isoforms. As shown in Figure 1H, BRAF mutants with a deletion of either Q494L495 or Q496A497 in the β3-αC loop stimulated the MEK-ERK pathway when expressed in 293T cells, and the central RH alteration (R509H) in the dimer interface abolished their activity. This suggests that BRAF can also be activated by the β3-αC loop deletion-driven homodimerization.

### ARAF has both allosteric and catalytic activities at lower levels than those of BRAF and CRAF

The fact that ARAF signaling through ERK was activated by Q347A348 deletion-driven homodimerization indicated that it is able to function as both allosteric activator and receiver. To confirm this, we carried out a set of RAF co-activation assays ^20, 36, 37^. In these assays, ARAF(AGFF) is a N-terminal truncated mutant (aa285-606) with a non-phosphorylatable NtA motif that functions as a receiver, whereas ARAF(DGEE/V324F) is a similar truncatant with an acidic NtA motif and a fused catalytic spine ^38, 39^ that mimics the inhibitor-bound status and works as a kinase-dead allosteric activator (Table S1). When co-expressed in 293T cells with allosteric activators derived from different RAF isoforms, ARAF(AGFF) was strongly activated by BRAF, intermediately by CRAF, and weakly by itself through dimerization (Figure 2A). On the other hand, ARAF(DGEE/V324F) stimulated moderately the catalytic activity of ARAF and CRAF receivers, but only slightly that of BRAF receiver when co-expressed in 293T cells (Figure 2B-C). Taken together, these data suggest that ARAF can function as both receiver and allosteric activator, albeit less efficiently than the other two isoforms.

**Figure 2.**
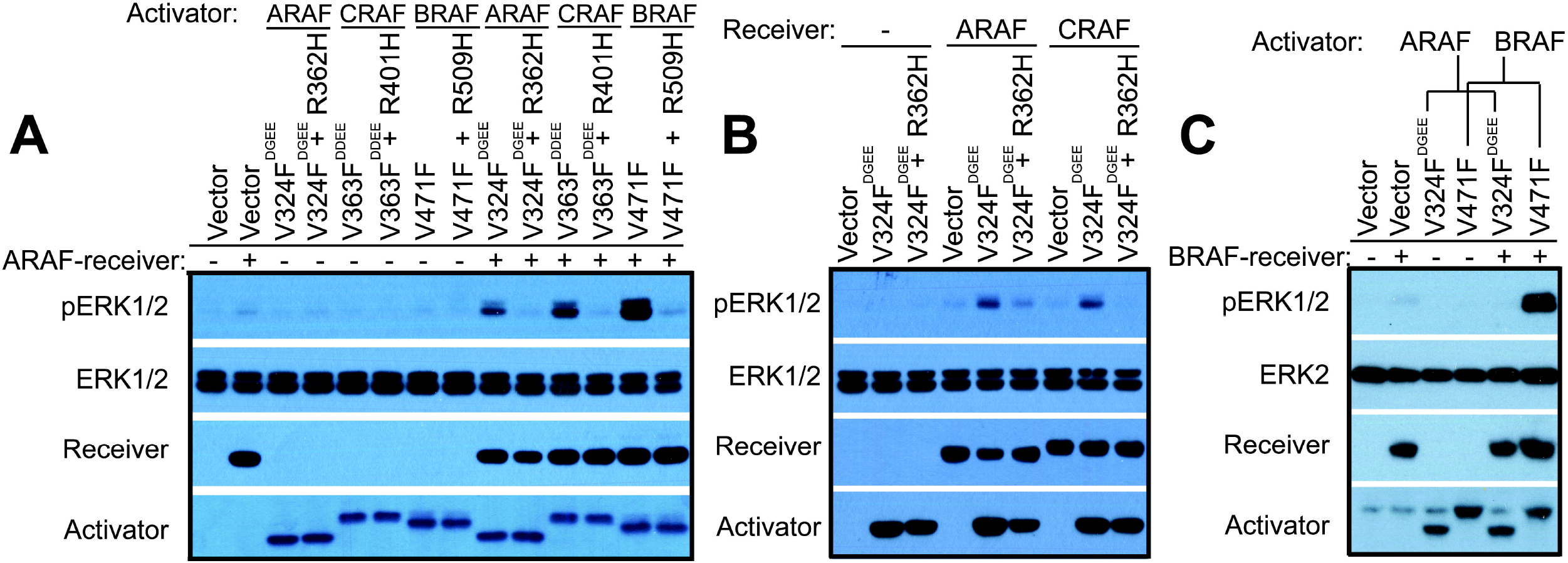
ARAF has both allosteric and catalytic activities albeit less than BRAF and CRAF. (A) As a receiver, ARAF is strongly activated by BRAF, intermediately by CRAF, and weakly by itself through dimerization. The RAF co-activation assays were carried out as before ^20, 37^. Briefly, the activator and the receiver were co-expressed in 293T cells, and the phospho-ERK1/2 was measured by immunoblot. (B-C) As an activator, ARAF stimulates moderately the catalytic activity of ARAF and CRAF receivers, albeit hardly that of BRAF receiver. The RAF co-activation assays were carried out as in (A). All images are representative of at least three independent experiments.

### The weak activity of ARAF arises from its non-canonical APE motif that decreases its dimer affinity

Regulatory spine (R-spine) is a hallmark signature of active protein kinases that comprised of four conserved residues, namely RS1-4 ^38, 39^. To explore the molecular basis underlying the weak activity of ARAF, we examined whether the R-spine-favored mutation could turn ARAF into a constitutively active kinase independent of dimerization-driven transactivation, as it has been shown for CRAF and BRAF ^20, 36, 38-41^. Similar to BRAF and CRAF, an ARAF mutant combining a RS3 mutation (L358M) and a negatively charged NtA motif showed a strong activity towards MEK-ERK pathway when expressed in 293T cells (Figure 3A). Although the activity of this mutant (ARAF, DGEE/L358M) did not depend on the AL (<u>a</u>ctivation <u>l</u>oop)-phosphorylation, it was impaired by the central RH alteration (R362H) in the dimer interface (Figure 3B), indicating again that ARAF has different characteristics from BRAF and CRAF. By aligning the kinase domain sequences of RAF isoforms, we found that ARAF had a non-canonical APE motif whose Pro is altered into Ala (Figure 3C). The APE motif localizes at the N-terminus of α-helix EF (αEF), and the Pro is a helix breaker that makes the N-terminus of αEF flexible. The Glu (E) next to Pro in the APE motif forms a salt-bridge with Arg in the αH-αI loop (Figure 3D), which has been shown to play a critical role in the regulation of protein kinase A ^42^. Since Ala is a helix-favored residue, we thought that a substitution of Pro with Ala would generate a longer αEF with a more rigid N-terminus, and therefore weaken/break the Glu-Arg salt-bridge and impair kinase function. To test this hypothesis, we mutated Ala back to Pro in the APE motif of ARAF(DGEE/L358M), and found that this mutant was resistant to the central RH alteration in dimer interface as well as the homologous BRAF and CRAF mutants described in our previous studies ^20, 37^ (Figure 3E). Furthermore, ARAF activator and receiver with a canonical APE motif exhibited much stronger activity than their wild-type counterparts in co-activation assays (Figure 3F&G). On the other hand, a substitution of Pro with Ala in the APE motif of BRAF and CRAF sensitized their constitutively-active R-spine mutants, BRAF(L505M) and CRAF(DDEE/L397M), to the central RH alteration in dimer interface (Figure 3H&I). Moreover, a breakage of the Glu-Arg salt-bridge by replacing Arg with Gln also led to the sensitivity of CRAF (DDEE/L397M) to the central RH alteration in dimer interface (Figure 3I). Together, these data demonstrate that the different APE motifs of RAF isoforms are responsible for the differential activities observed between ARAF and the other RAF paralogs.

**Figure 3.**
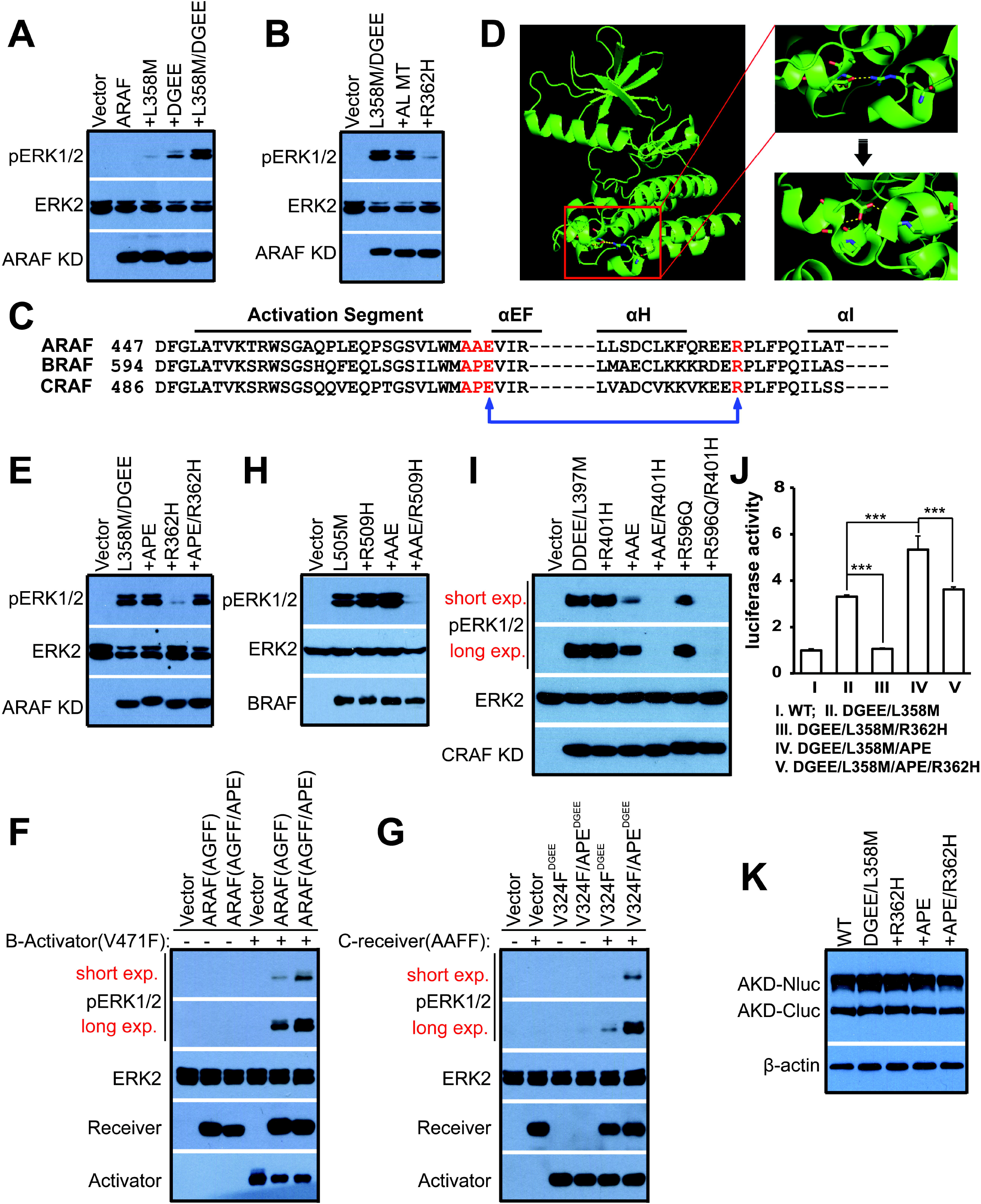
The non-canonical APE motif decreases the activity of ARAF. (A) R-spine fusion together with acidic NtA motif fully activates ARAF. (B) The constitutively-active R-spine mutant of ARAF is resistant to the activation loop (AL) mutation but not to the central RH alteration in dimer interface. (C) ARAF has a non-canonical APE motif. The conserved APE motif is altered into AAE in ARAF, which might weaken the Glu-Arg salt bridge between APE motif and αH-αI loop. (D) Schematic diagram showing the Glu-Arg slat bridge in CRAF. Schematic diagram of CRAF (PDB ID: 3OMV) was generated by using PyMOL software. (E) The conserved APE motif restores the activity of ARAF R-spine mutant with the central RH alteration in dimer interface. (F-G) The conserved APE motif enhances both allosteric activator and receiver activities of ARAF. The RAF co-activation assays were carried out as in Figure 2. (H) The alteration of APE motif makes BRAF R-spine mutant sensitive to the central RH alteration in dimer interface. (I) The alteration of APE motif or the breakage of Glu-Arg salt bridge makes CRAF R-spine mutant sensitive to the central RH alteration in dimer interface. (J-K) The conserved APE motif enhances the dimer affinity of ARAF mutants. The dimer affinity of ARAF mutants is measured by complementary split luciferase as in Figure 1A (*n*=5, \*\*\**p*<0.001). In (A-B), (E), and (H-I) the activity of RAF mutants in 293T transfectants was measured by anti-phospho-ERK1/2 immunoblot. All images are representative of at least three independent experiments. “KD” represents for “kinase domain” in the full text.

In order to further address how the non-canonical APE motif of ARAF dampens its functional activities, we next investigated whether it affects the dimer affinity of ARAF by using complementary split luciferase assay as described above. As shown in Figure 3J&K, a canonical APE motif conferred the higher dimer affinity of ARAF R-spine mutants, comparing ARAF(DDEE/L358M) with ARAF(DDEE/L358M/APE), and ARAF(DDEE/L358M/R362H) with ARAF(DDEE/L358M/R362H/APE), which indicates that the non-canonical APE motif of ARAF decreases its dimer affinity. Since the dimerization plays a critical role in RAF activation, this data indicates that the lower dimer affinity arising from the non-canonical APE motif leads to the weaker activities of ARAF among RAF isoforms. In addition, the lower dimer affinity of ARAF(DDEE/L358M) and its sensitivity to the central RH alteration in dimer interface in contrast to ARAF(DDEE/L358M/APE), suggest that the active R-spine mutants of ARAF might function as a dimer to activate MEK-ERK pathway even if they do not require dimerization-driven transactivation for triggering their activity.

### Similar in-frame deletions of dimeric protein kinases exist in cancer genomes

Dimerization-driven allostery plays a key role not only in the activation of RAF kinase but also in that of many other protein kinases ^43, 44^. Besides ARAF(ΔQA), an oncogenic BRAF mutant with β3-αC loop deletion (ΔNVTAP) has been reported although its activation mechanism remains controversial ^31-34^. We here aimed to explore all kinase mutants with similar deletions in cancer genomes and to asses the importance of these mutations in human cancers. To this end, we interrogated the ICGC (International Cancer Genome Consortium) database, the cBioportal for Cancer Genomics database, and the COSMIC (Catalogue of Somatic Mutations in Cancer) database, and summarized all kinase mutants with similar in-frame deletions of β3-αC loop, including those reported in literatures ^31-34^, in Table S2. Among these mutants, the EGFR exon19 del is highly prevalent and has been shown to elevate kinase activity by promoting side-to-side homodimerization ^45^. Other similar kinase mutants include those of BRAF, JAK1 and ERB B2, which (except JAK1 mutants, which have not been tested) have shown elevated kinase activity (see below and Figure S2).

### In-frame deletions of β3-αC loop activate BRAF as well as CRAF by enforcing homo-dimerization

To characterize BRAF mutants in Table S2 and determine whether they are activated by enhanced homodimerization, we expressed these mutants in 293T cells and fibroblasts. All these mutants stimulated the MEK-ERK pathway independent of upstream stimuli or endogenous RAF molecules (Figure 4A&B and S3A), indicating that they are constitutively active. However, these mutants exhibited differential resistance to the central RH alteration (R509H) in dimer interface (Figure 4C). This alteration had no effect on the activity of BRAF(ΔNVTAP), partially inhibited that of BRAF(ΔMLN), and completely blocked that of BRAF(ΔNVTAPT). We reasoned that this discrepancy among BRAF mutants might arise from their different dimer affinity/stability. To test this hypothesis, we carried out co-immunoprecipitation assays, and found that indeed these mutants had enhanced but different propensities to form dimers, with ΔNVTAP > ΔMLN > ΔNVTAPT ≈ ΔQA > WT, independently of ERK1/2-mediated feedback ^46^ (Figure 4D and S3B&C). The R509H alteration prevented the homodimerization of BRAF(ΔNVTAPT) and BRAF(ΔQA), partially that of BRAF(ΔMLN), and weakly that of BRAF(ΔNVTAP). Previous studies have shown that although the central Arginine alteration impairs the dimerization-driven transactivation of wild-type RAFs ^15^, it makes up less than 20% dimer interface ^11^. Therefore, the resistance of BRAF(ΔMLN) and BRAF(ΔNVTAP) to the central R509H alteration in dimer interface does not exclude that these two BRAF mutants are activated through dimerization-driven transactivation by virtue of their much stronger dimer affinity than wild-type counterpart. To further demonstrate that these BRAF mutants, especially BRAF(ΔNVTAP) and BRAF(ΔMLN), are activated by enhanced homodimerization, we performed the RAF co-activation assay ^20, 36, 37, 40, 41^ by using their kinase-dead V471F mutants as allosteric activators. All activators tested in this study strongly stimulated the catalytic activity of CRAF receivers (Figure 4E), and particularly activators derived from BRAF(ΔNVTAP) and BRAF(ΔMLN) could even trigger endogenous RAF molecules (Figure 4E lane5 & lane3). The central R509H alteration in dimer interface was unable to prevent these two strong allosteric activators from triggering BRAF receivers (Figure S3D). Since the non-canonical APE motif had been also shown to decrease the dimer affinity in RAF molecules, we next introduced it together with the central R509H alteration into BRAF(ΔNVTAP) mutant, and found that this combined alteration completely blocked the activity of BRAF(ΔNVTAP) (Figure S3E). Taken together, our data demonstrates that all BRAF mutants with in-frame deletions of β3-αC loop are activated by enhanced homodimerization.

**Figure 4.**
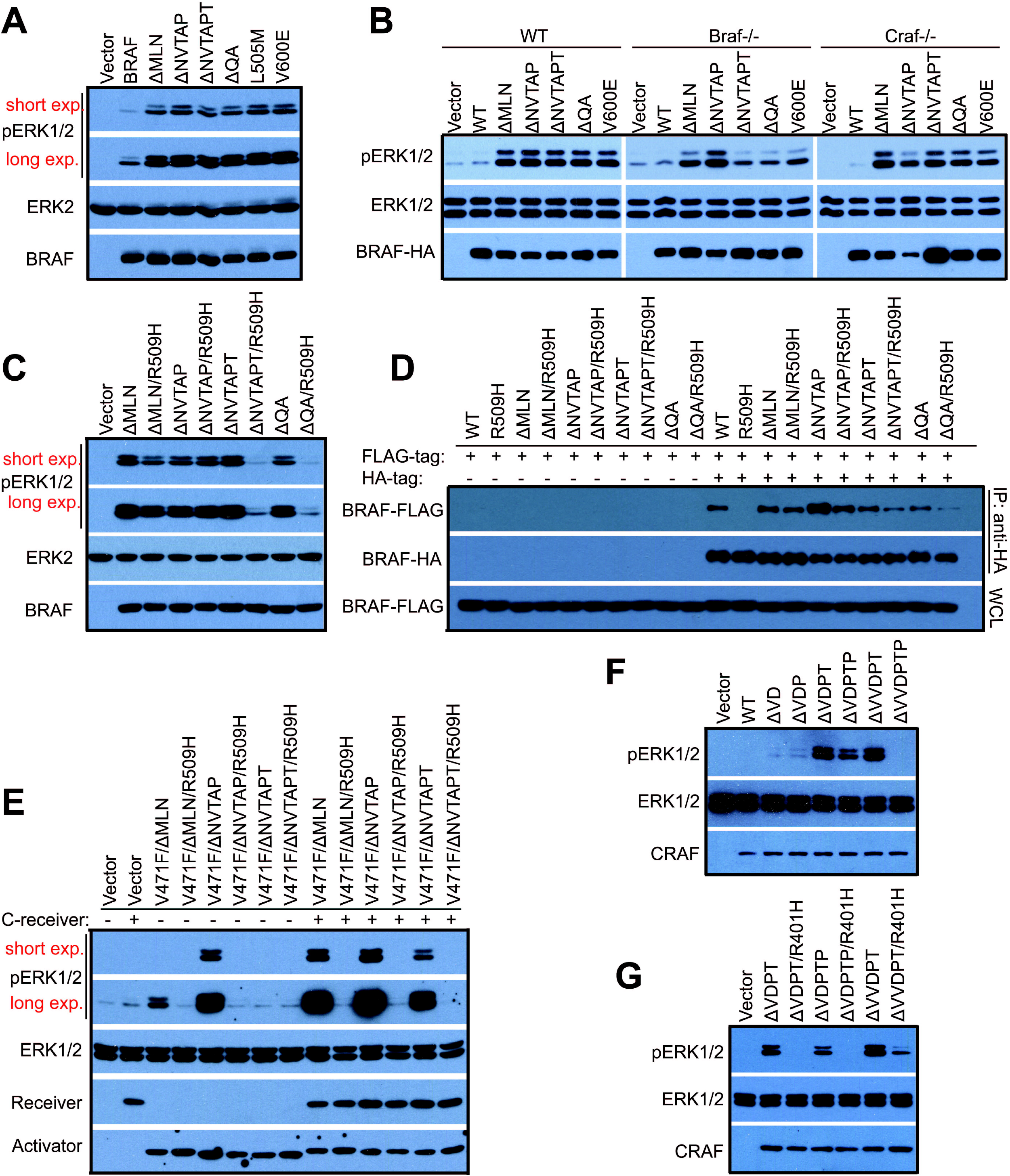
In-frame β3-αC loop deletions activate RAF kinase by enhancing homodimerization. (A-C) BRAF mutants with in-frame β3-αC loop deletions activate the MEK-ERK pathway independent of endogenous RAF molecules, and exhibit distinct resistance to the central R509H alteration in dimer interface. The activity of BRAF mutants in 293T transfectants or MEF stable cell lines was measured by anti-phospho-ERK1/2 immunoblot. (D) BRAF mutants with in-frame β3-αC loop deletions have an elevated but differential dimer affinity/stability. BRAF mutants with FLAG- or HA-tag were coexpressed in 293T cells, and immunoprecipitated by anti-HA beads and detected by anti-FLAG immunoblot. (E) BRAF mutants with in-frame β3-αC loop deletions have a strong allosteric activity. Catalytic spine-fused BRAF mutants were expressed with or without CRAF-receiver in 293T cells, and their ability to stimulate the downstream pathway was measured as phospho-ERK1/2 in 293T transfectants. (F-G) The β3-αC loop deletions activate CRAF by enhancing homodimerization. CRAF mutants were expressed in 293T cells, and their activity was measured as phospho-ERK1/2. All images are representative of at least three independent experiments.

Unlike ARAF and BRAF, we did not find any CRAF mutants with in-frame deletions of β3-αC loop in databases. To test whether CRAF can be activated by this type of mutations, we constructed mutants homologous to those of ARAF and BRAF. When expressed in 293T cells, ΔVDPT, ΔVDPTP, and ΔVVDPT mutants of CRAF, but not other mutants, strongly activated the MEK-ERK pathway independent of upstream stimuli (Figure 4F and S3F&G), and exhibited differential resistance to the central RH alteration (R401H) in dimer interface (Figure 4G). Since we had showed that BRAF was activated by mutations (ΔQA and ΔQL) homologous to ARAF(ΔQA), we hence determined whether ARAF could be triggered by mutations homologous to BRAF(ΔMLN, ΔNVTAP, ΔNVTAPT). However, none of these alterations activated ARAF (Figure S3H).

### All BRAF mutants with variable deletions of β3-αC loop have a strong transforming potential, and a robust resistance to Vemurafenib but not LY3009120 that correlates with their dimer affinity

Although the oncogenic potential and resistance to RAF inhibitor of BRAF(ΔNVTAP) has been reported recently ^31, 34^, whether all BRAF mutants with in-frame deletions of β3-αC loop are able to function as cancer drivers and their pharmacological characteristics are not clear. To address these questions, we first measured the oncogenic potential of these mutants by foci formation assays. As shown in Figure 5A and S4A&B, all BRAF mutants with in-frame deletions of β3-αC loop transformed immortalized fibroblasts and induced foci formation independent of endogenous RAF molecules, suggesting that they can function as drivers to induce cancers. Further, we examined their sensitivities to the RAF inhibitor Vemurafenib, and found that all mutants exhibited a robust resistance to this drug, ranking as BRAF(ΔNVTAP) > BRAF(ΔMLN) > BRAF(ΔNVTAPT) ≈ BRAF(ΔQA) >> BRAF(V600E) (Figure 5B&C), which correlates with their dimer affinity/stability. However, all these mutants had similar sensitivities to the RAF dimer inhibitor, LY3009120, which are comparable with that of BRAF(V600E) in A101D melanoma cell line (Figure 5D&E).

**Figure 5.**
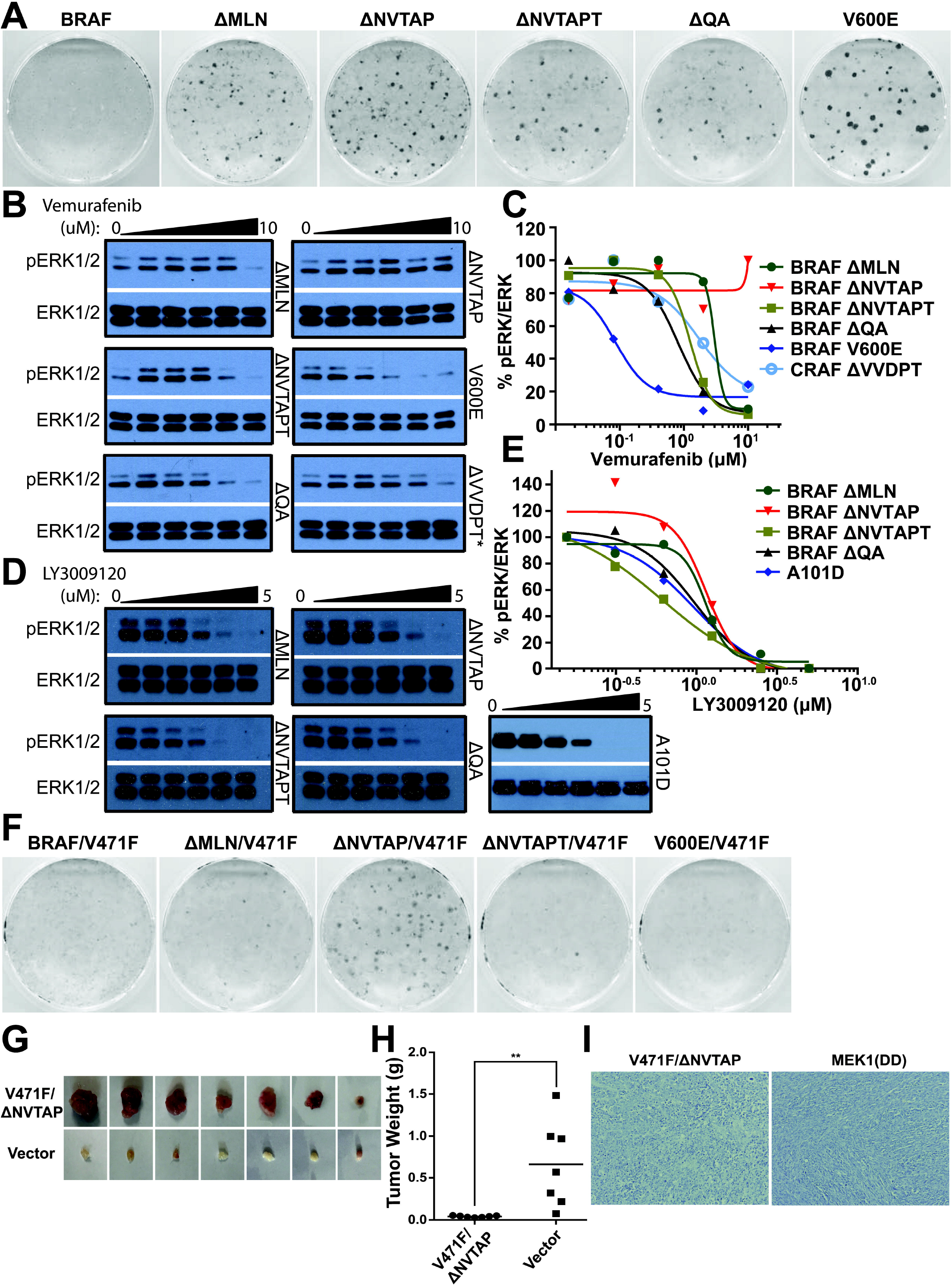
BRAF mutants with in-frame β3-αC loop deletions have a strong transforming ability and a robust but differential resistance to Vemarufenib. (A) BRAF mutants with in-frame β3-αC loop deletions induce foci formation when stably expressed in immortalized fibroblasts. The foci formation assay was carried out as in Figure 1C. (B-C) BRAF mutants with in-frame β3-αC loop deletions exhibit a robust but differential inhibitor resistance. Stable fibroblast cells that express individual BRAF mutants with in-frame β3-αC loop deletions were treated with Vemurafenib for 4 hours, and phospho-ERK1/2 was probed by immunoblot and quantified by using Image J. The graphs were generated by using GraphPad Prism 6. (D-E) BRAF mutants with in-frame β3-αC loop deletions have approximate sensitivities to RAF dimer inhibitor, LY3009120. The drug sensitivities of BRAF mutants were measured as in (B-C). The BRAF(V600E)-harboring melanoma cell line, A101D, was used as control. (D-G) Catalytic spine-fused BRAF mutants with in-frame β3-αC loop deletions have variable oncogenic potentials *in vitro* and *in vivo*. D, the oncogenic potential of BRAF mutants was measured by the foci formation assay as in (A). E, Xenograft tumors were generated in NOD/SCID mice from immortalized fibroblasts that express BRAF mutants as described in Materials & Methods. F, the weight of xenogrfat tumors from E (*n*=7 for each group, \*\**p*<0.01). G, Representative images from histological section staining of xenograft tumors from E (*n*=7). The MEK1DD-driven xenograft tumor served as control. All images are representative of at least three independent experiments.

### The high dimer affinity of kinase-dead BRAF mutants with β3-αC loop deletions bypasses the requirement of active RAS to drive tumorigenesis

The inhibitor-loading has been shown to fuse the catalytic spine of RAF molecules, which can be mimicked by the Val to Phe mutation in ATP-binding pocket ^20, 36-41^. Compound BRAF mutants with both β3-αC loop deletion and Val471Phe, especially BRAF(ΔNVTAP/V471F), lacked catalytic activity but were able to activate the MEK-ERK pathway through triggering wild-type RAFs (Fig4E). The kinase-dead BRAF(ΔNVTAP/V471F) induced foci formation *in vitro* and tumor formation *in vivo* even in the absence of active RAS, but dependent on endogenous RAF molecules (Figure 5F-I and S4C). Since a previous study had shown that kinase-dead RAFs cooperate with active RAS to induce tumorigenesis ^12^, this data suggests that the high dimer affinity of RAF mutants could bypass the requirement of active RAS to driven cancer development.

### Active RAF kinases function as a dimer to phosphorylate MEK

The activation of the RAF/MEK/ERK kinase cascade is a very complex process and its underlying molecular basis is not completely understood ^45-49^. In current model, RAF and MEK form a face-to-face dimer in quiescent cells. Upon stimulation, two RAF-MEK dimers are brought together and assemble a transient tetramer through back-to-back RAF dimerization, which not only activates RAF but also facilitates subsequent MEK activation ^19^. However, how active RAF activates MEK is not clear. To elucidate this process, we first tested whether the dimerization of active RAF is required for MEK activation by using BRAF mutants with in-frame deletions of β3-αC loop since these mutants have different dimer affinity/stability. As shown in Figure 6A, those constitutively-active BRAF mutants with low dimer affinity/stability such as BRAF(ΔNVTAPT) and BRAF(ΔQA) lost their catalytic activity towards MEK *in vitro* when purified by immunoprecipitation, in contrast to those mutants with high dimer affinity/stability which retained catalytic activity toward MEK. We reasoned that the loss of activity of BRAF mutants *in vitro* arises from their dimer dissociation during purification. To test this hypothesis, we strengthened the dimers of BRAF(ΔNVTAPT) and BRAF(ΔQA), respectively, by GST fusions ^50^, and found that it restored their catalytic activity *in vitro* (Figure 6B). This phenomenon was also seen with a homologous ARAF mutant (ΔQA) whose *in vitro* catalytic activity was rescued by GST fusion (Figure 6C). As reported before ^20, 51^ and shown in this study, both alterations of the central RH in dimer interface and the APE motif significantly impair but do not completely abolish the dimer formation of RAF molecules. We therefore next examined the effect of these alterations on the *in vitro* catalytic activity of active RAF mutants. Among three active ARAF R-spine mutants, only the one with a high dimer affinity (see Figure 3J-K) maintained its catalytic activity *in vitro* after purification (Figure 6D), and GST fusion restored that of the other two mutants with a low dimer affinity (Figure 6E). Similar to that of ARAF mutants, active CRAF mutants with an alteration of RH, or of APE, lost their catalytic activity *in vitro*, which was recovered by GST fusion-enhanced dimerization (Figure 6F). As to active BRAF R-spine mutants, the alterations of R509H, or APE, inhibited their catalytic activity *in vitro* by different extends, which was also relieved by the GST fusion approach (Figure 6G). The loss of in vitro catalytic activity of RAF mutants with low dimer affinity could be also rescued by other dimeric molecular fusions (data not shown), or partially restored with a gentle wash of PBS during purification (Figure S5), which excludes the potential artificial effect arising from GST fusion. Together, these data indicate that all RAF isoforms would function as dimers to phosphorylate MEK.

**Figure 6.**
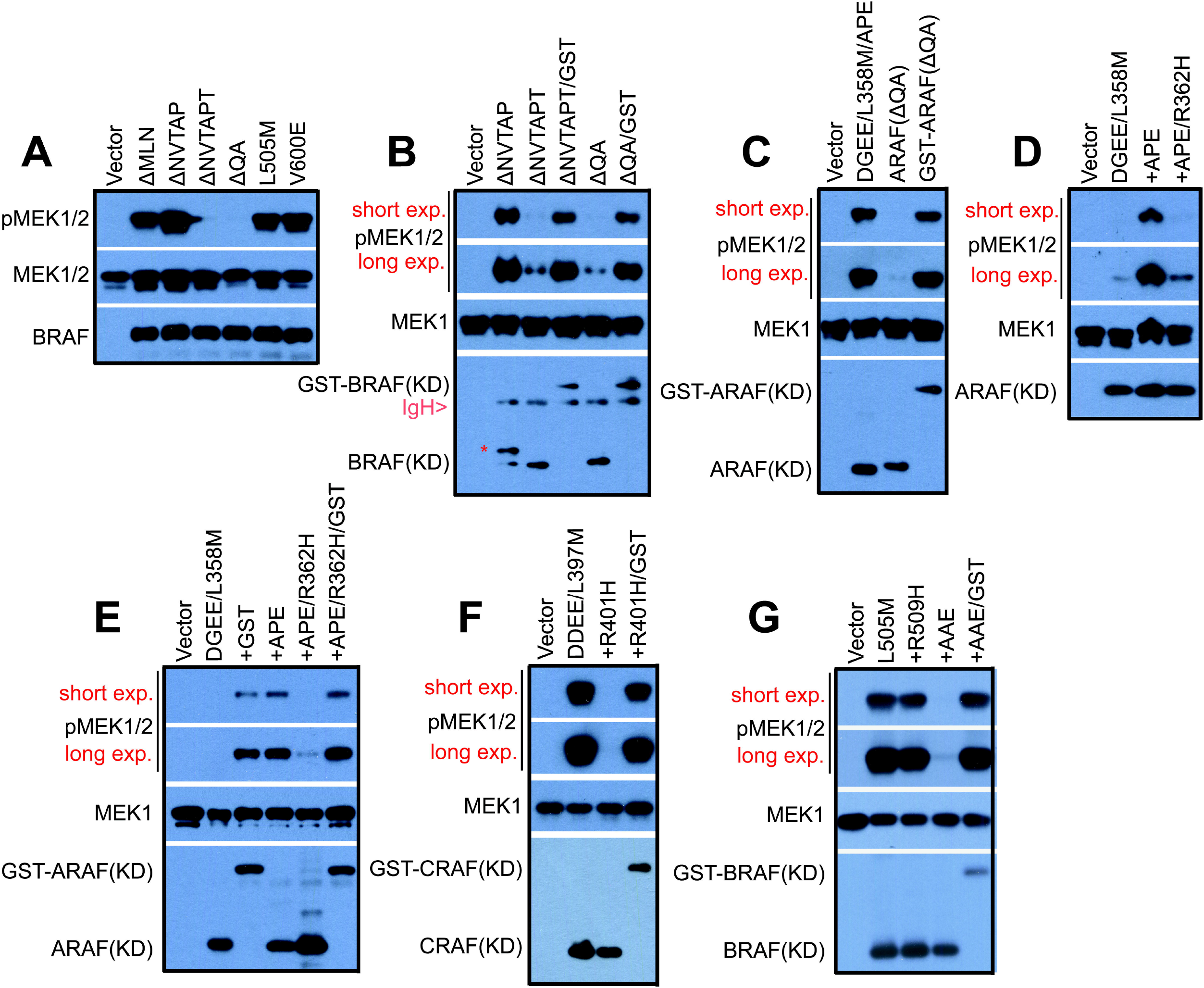
Active RAF mutants phosphorylate MEK in a dimer-dependent manner. (A-B) BRAF mutants with in-frame β3-αC loop deletions lose their catalytic activity in vitro upon purification if they have a low dimer affinity, which is rescued by GST fusion. The band labeled with “*” in lane 2 of B represents the highly phosphorylated BRAF(KD, ΔNVTAPT). (C) Like BRAF(ΔQA), purified ARAF(ΔQA) loses its kinase activity in vitro, which is rescued by GST fusion. (D-G) GST fusion restores in vitro the kinase activity of constitutively active R-spine mutants of ARAF, CRAF, and BRAF with low dimer affinity. In A-G, all RAF mutants were expressed in 293T cells and purified by immunoprecipitation, and their activity was measured by in vitro kinase assays as described before ^20, 37^ All images are representative of at least three independent experiments.

### BRAF(V600E) also phosphorylates MEK through a dimer-dependent manner

Given its resistance to the central RH alteration in dimer interface and sensitivity to Vemurafenib, BRAF(V600E), a dominant mutant in RAF mutation spectrum, has been proposed to function as a monomer to activate MEK ^26^. However, recent studies showed that BRAF(V600E) exists as dimers or high-order oligomers in cells ^51-53^. This prompted us to determine whether, unlike other active RAF mutants, BRAF(V600E) truly phosphorylates MEK in a dimer-independent manner. Since BRAF(V600E) has an enhanced propensity to form dimers ^51-53^, the central RH alteration (R509H) in dimer interface is unable to dissociate its dimers completely. Hence, our approach was to replace one protomer in the dimer of BRAF(V600E) with one dysfunctional BRAF mutant (loss of both catalytic and allosteric activities), and measured the catalytic activity of BRAF(V600E) in heterodimers. To generate such a dysfunctional BRAF mutant, we mutated the residues of the kinase-dead BRAF(ΔNVTAP/V471F) that mediate the heterodimerization of BRAF with MEK ^19^ (Figure 7A). Since BRAF utilizes two different groups of residues to bind MEK and RAF ^19^, the introduction of these mutations would not influence RAF dimerization. Unlike its prototype, the mutant, BRAF(ΔNVTAP/V471F/R462E/I617R/F667A), heretofore referred to as BRAF(ΔNVTAP/V471F)*, had no allosteric ability to trigger endogenous RAF molecules when expressed in 293T cells (Figure 7B). Moreover, BRAF(V600E) bound to BRAF(ΔNVTAP/V471F)* had little catalytic activity *in vitro*, in contrast to that bound to BRAF(ΔNVTAP/V471F) when purified from 293T co-transfectants (Figure 7C). This suggests that a BRAF(V600E) molecule needs a partner able to hold MEK in order to phosphorylate it. To further confirm this finding, we introduced BRAF(ΔNVTAP/V471F)* into BRAF(V600E)-dependent melanoma cell lines by lentiviral transductions and found that its expression down-regulated phospho-ERK1/2 and inhibited cell growth in vitro and xenograft tumor growth in vivo (Figure 7D-H).

**Figure 7.**
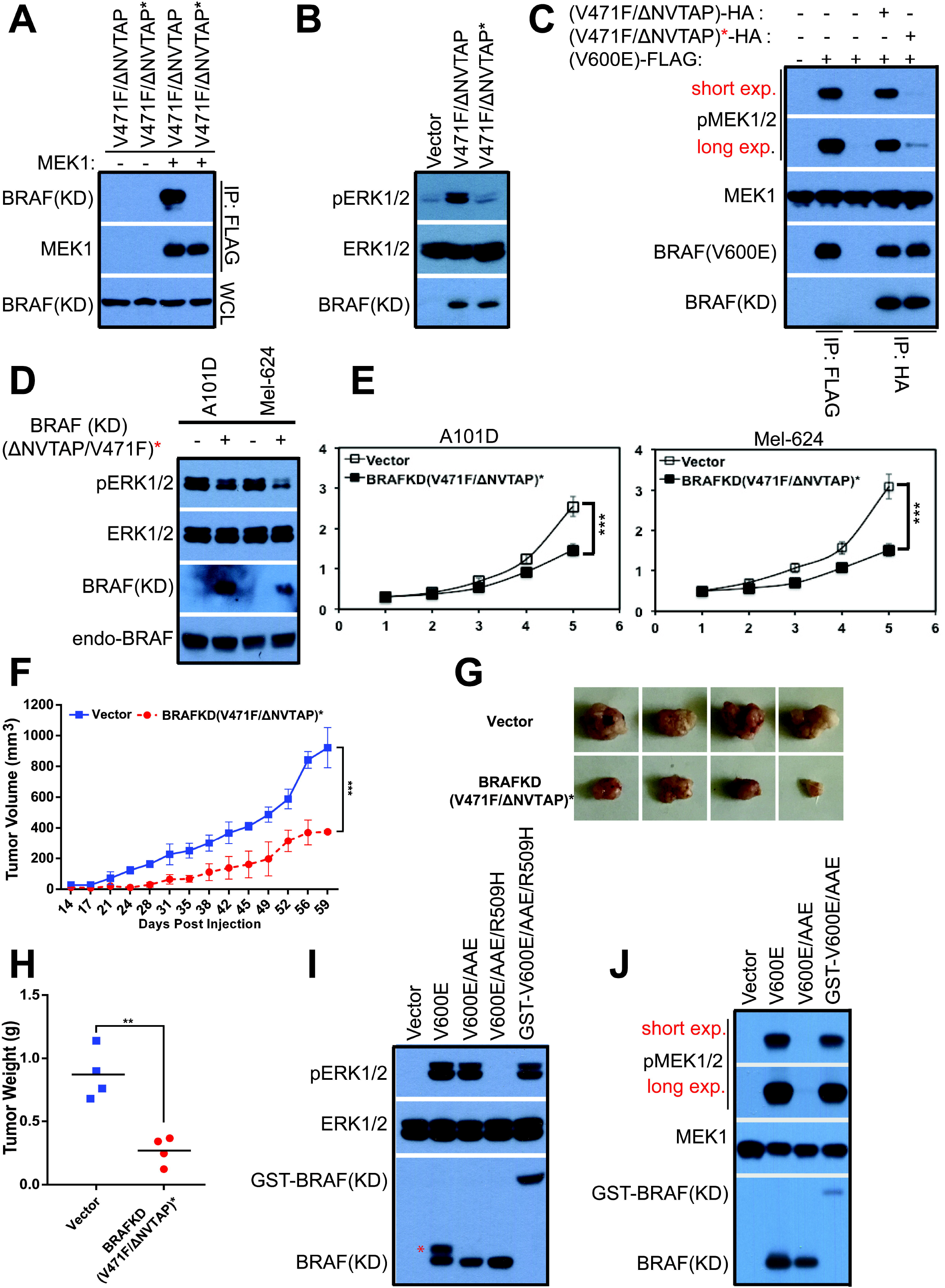
BRAF(V600E) functions as a dimer to activate MEK. (A-B) The kinase-dead BRAF mutant with high RAF dimer affinity requires ability to heterodimerize with MEK1 for transactivating wild-type RAF molecules. A, BRAF(ΔNVTAPT/V471F/R462E/I617R/ F667A), referred to as BRAF(ΔNVTAPT/V471F)* below, was generated by PCR and its ability to heterodimerize with MEK1 was measured by co-immunoprecipitation as in Figure 1E. B, Unlike its prototype, BRAF(ΔNVTAPT/V471F)* is not able to activate endogenous RAF molecules when expressed in 293T cells. BRAF mutants were expressed in 293T cells, and their activity was measured by anti-phospho-ERK1/2 immunoblot. (C) BRAF(V600E) loses its catalytic activity once dimerizing with BRAF(ΔNVTAPT/V471F)*. BRAF(V600E) that binds to BRAF(ΔNVTAPT/V471F)* was purified by immunoprecipitation from 293T cotransfectants and its activity was measured by in vitro kinase assay as in Figure 6. BRAF(V600E) that binds to BRAF(ΔNVTAPT/V471F) was expressed and purified from 293T cotransfectants, and serviced as a control. (D-H) The stable expression of BRAF(ΔNVTAPT/V471F)* in BRAF(V600E)-dependent melanoma cell lines dampens MEK-ERK signaling, and inhibits cell growth in vitro and xenograft tumor growth in vivo. D-E, BRAF(ΔNVTAPT/V471F)* was stably expressed in A101D and Mel-624 cell lines, and phospho-ERK1/2 and cell growth were measured respectively by immunoblot or by cell counting (*n*=5, \*\*\**p*<0.001). F-H, the growth curve, the photos and the weight of xenograft tumors derived from A101D melanoma cell lines that stably express BRAF(ΔNVTAPT/V471F)* or empty vector (*n*=4 for each group, \*\*\**p*<0.001, \*\**p*<0.01). (I-J) The activity of BRAF(V600E) with non-canonic APE motif is abolished by the central RH alteration in dimer interface in vivo or upon purification in vitro by immunoprecipitation, which is restored by GST fusion. I, the APE alteration of BRAF(V600E) has little effect on its activity in cells, but makes BRAF(V600E) sensitive to the central RH mutation in dimer interface. BRAF mutants were expressed in 293T cells and their activity was measured by anti-phospho-ERK1/2 immunoblot. * in lane 2 indicates a post-translational modified version of BRAF(V600E). J, the BRAF(V600E) with non-canonic APE loses its catalytic activity upon immunoprecipitation, which can be rescued by GST fusion. BRAF mutants were expressed in 293T cells and purified by immunoprecipitation, and their activity was measured by in vitro kinase assay as in Figure 6. All images are representative of at least three independent experiments.

As shown above, the APE motif of RAF kinases regulates their dimerization, likely through affecting the Glu-Arg salt bridge between the APE motif and the αH-αI loop, and the alteration of APE motif into non-canonical AAE dramatically decreases RAF dimer affinity/stability. Therefore, we next determined whether an alteration of APE motif into AAE would impair the catalytic activity of BRAF(V600E) in vitro, as which occurs in ARAF R-spine mutants. As shown in Figure 7I-J and S6A, although the AAE variant of BRAF(V600E) had comparable ability with its prototype to phosphorylate MEK and thus turn on ERK signaling when expressed in 293T cells, it lost its catalytic activity upon purification by immunoprecipitation, which was restored by GST fusion. Moreover, the combined alterations of AAE and R509H completely blocked the activity of BRAF(V600E) even in vivo, which was also recovered by GST fusion (Figure 7I). To directly confirm that the discrepant catalytic activity among BRAF(V600E), BRAF(V600E/AAE), and BRAF(V600E/AAE/R509H), arises from their different dimer affinity/stability, we completed a co-immonprecipitation assay with a gentle wash of PBS as in Figure S5, and found that these BRAF variants had quite different ability to form dimers as BRAF(V600E) >> BRAF(V600E/AAE) >> BRAF(V600E/AAE/R509H) (Figure S6B). Altogether, these data demonstrate that like other RAF molecules, BRAF(V600E) also functions as a dimer to activate MEK.

## Discussion

The dimerization of RAF kinase not only plays a critical role in the activation of the RAF/MEK/ERK kinase cascade ^9-17^, but also contributes to drug resistance in cancer therapy ^11, 12, 14, 26^. Previous studies have shown that the RAF dimerization could be improved by active RAS, inhibitor binding, gene fusions or alternative splicing ^9, 11-14, 17, 22-26, 54^. However, whether it can be achieved by other ways remains unknown. Recently, some oncogenic RAF mutants with β3-αC loop deletions have been reported by several groups^31-34^, and both dimer-dependent and –independent models have been suggested to explain how this type of mutations activates RAF^32, 34^. Chen et al characterized BRAF(ΔLNVTAP) that resembles BRAF(ΔNVTAPT) in cancer genomes, and found that its activity was blocked by the central R509H alteration in dimer interface^32^. On the other hand, Foster et al showed that the other cancer-related mutant, BRAF(ΔNVTAP) was resistant to the same alteration^34^. Both groups failed to understand the discrepancy among these RAF mutants. In this study, we systemically characterize all RAF mutants with β3-αC loop deletions, and provide solid evidence that this type of mutations activates RAFs through improving homodimerization, which clarifies the controversial between those two groups.

Among RAF isoforms, BRAF and CRAF have been shown to function as both catalysis-competitive kinases and allosteric activators ^21^, while ARAF has a bare activity and been thought as a scaffold ^55, 56^. By characterizing the ARAF mutant with a β3-αC loop deletion (ΔQA), we demonstrate that like the other two paralogs, ARAF has both catalytic and allosteric activities though less. Further, we elucidate that the weak activity of ARAF arises from its non-canonical APE motif that decreases its dimer propensity likely by weakening the Glu-Arg salt bridge between the APE motif and the αI-αH loop. Our data clearly explain why ARAF has less ability to stimulate downstream MEK-ERK signaling.

In-frame deletions of β3-αC loop activate not only RAFs but also other dimeric protein kinases by enhancing homodimerization. Remarkably, we found this type of mutation in BRAF, ERBB2, JAK2, and EGFR in cancer genomes, suggesting that the dimerization works as a principal mechanism to regulate the activity of these kinases. All BRAF mutants with variable deletions of β3-αC loop exhibit the elevated albeit differential dimer affinity, which is reflected by their different resistance to the central R509H alteration in dimer interface. The strong dimer affinity of BRAF(ΔNVTAP) bypasses the requirement of active RAS for its kinase-dead version to activate endogenous RAF and transform cells, suggesting that the dimerization of RAF is a dynamic equilibrium in cells and active RAS shifts this equilibrium in the favor of dimers.

A strong dimerization is required for transactivation of RAF molecules, whereas a weak dimerization is still indispensable for their catalytic function. The previous conclusion that active RAF kinase may function as a monomeric enzyme is mainly based on the data from the central RH alteration in dimer interface of RAF molecules. Recent studies revealed that this alteration was not able to completely dissociate RAF dimers although it blocked the dimerization-driven transactivation of RAF molecules ^20, 51^. Here, we found that this mutation does not block the dimerization-driven transactivation RAF mutants with high dimer affinity, such as BRAF(ΔNVTAP) and BRAF(ΔMLN), which calls for a re-evaluation of previous data resulting from the central RH alteration in dimer interface. Constitutively active RAF mutants including BRAF(V600E) activate MEK in a dimer-dependent manner even if they do not require a dimerization to trigger their activity, implying that RAF kinase serves as both an enzyme and a MEK-docking platform in its catalysis process. This is further supported by that the kinase-dead BRAF mutant with high dimer affinity is not able to activate downstream MEK-ERK pathway through wild-type RAF molecules if it cannot bind MEK (Fig7A-B). These findings suggest that the MEK-docking platform function of RAF kinase can be used as a target to develop novel therapeutic inhibitors against both RAS- and RAF-driven cancers. In current cancer therapies, the efficacy of RAF and MEK inhibitors has been severely limited by intrinsic and acquired resistances arising from the paradoxical activation or re-activation of RAF/MEK/ERK kinase cascade ^57^. Such a new class of docking inhibitor would cover all type of cancers driven by hyperactive RAS/RAF/MEK/ERK signaling, and thus potentially have a greater and longer efficacy. Our study sheds a light on the treatment of RAS/RAF-driven cancers and has important clinic implications.

## Materials and Methods

### Biochemicals

Antibodies used in this study included: anti-phosphoERK1/2 (#4370), anti-phosphoMEK1/2 (#9154), and anti-MEK1/2 (#9124) (Cell Signaling Technology); anti-BRAF (SAB5300503), anti-CRAF (SAB5300393), anti-FLAG (F3165) and anti-β-actin (A2228) (Sigma); anti-HA (MAB6875, Novus Biologicals); anti-ERK2 (sc-154, Santa Cruz Biotechnology); anti-ERK1/2 (A0229, AB clonal); anti-Ki67 (ab16667, Abcam); and HRP-labeled secondary antibodies (Jackson Laboratories). Vemurafenib and LY3009120 were purchased from Medchemexpress; and D-luciferin from Gold Biotechnology. All other chemicals were obtained from Sigma.

### Plasmids and Cell lines

cDNAs encoding proteins in this study were purchased from Addgene or synthesized by Integrated DNA Technologies. All mutations were generated by PCR and tagged with either FLAG or HA or His, and cloned into vectors by Gibson assembly. pCDNA3.1(+) vector (Invitrogen) was used for transient expression; viral vectors (Clontech) for stable expression; and pET-28a (Novagen) for bacterial expression.

Wild-type, BRAF^-/-^ and CRAF^-/-^ MEFs were generated in previous study ^58, 59^. Melanoma cell lines: MeWo, A101D, Mel-624 were obtained from ATCC.

### Protein expression and Purification

6xhis-tagged MEK1 (K97A) and 6xhis-tagged ERK2(K52A) were expressed in BL21(DE3) strains and purified by using Nickel column (Qiagen) and following our previous protocol ^60^.

### Cell Culture, Transfection, and Transduction

All cell lines were maintained in DMEM medium with 10% FBS (Hyclone). Cell transfection were carried out by using the Biotool transfection reagent and following the manufacturer’s protocol. To generate stable cell lines that express RAF or MEK1 mutants, viruses were prepared and applied to infect target cells according to our previous studies ^20, 36, 37, 60, 61^. Infected cells were sorted by FACS or selected by using antibiotics.

### Immunoprecipitation, *In Vitro* Kinase Assay, and Western Blotting

Immunoprecipitations were performed as described previously ^20, 36, 37^. Briefly, whole-cell lysates were mixed with either anti-HA (E6779), or anti-FLAG beads (A2220) (Sigma), rotated in cold room for 60 min, and washed three times with RIPA buffer. For *in vitro* kinase assays, the immunoprecipitants were washed once with kinase reaction buffer (25 mM HEPES, 10 mM MgCl2, 0.5 mM Na3VO4, 0.5 mM DTT, pH 7.4), then incubated with 20μl kinase reaction mixture (2 ug substrate and 100 mM ATP in 20μl kinase reaction buffer) per sample at room temperature for 10 min. Kinase reaction was stopped by adding 5μl per sample 5XLaemmli sample buffer. Immunoblotting was carried out as described before ^20, 36, 37, 60, 61^.

### Foci formation assay

Immortalized MEFs infected with retroviruses encoding target proteins were plated at 5×10^3^ cells per 60mm dish, and fed every other day. 12 days later, cells were fixed with 2% formaldehyde and stained with Giemsa solution (Sigma).

### Complementary Split Luciferase Assay

293T transfectants that express different pairs of Nluc- and Cluc-fused RAF proteins were plated in 24-well Krystal black image plates at the the seeding density of 2×10^5^ per well. 24 hour later, D-luciferin (0.2mg/ml) with or without Vemurafenib (10μM) was added to the culture, and the incubation was allowed for 30 min before the luciferase signals were measured by Promega GloMax®-Multi Detection System.

### Animal studies

For xenograft experiments, female NOD/SCID mice (6~8 weeks) were injected with 3×10^6^ cells per mice in 1:1 matrigel (Corning). Tumor volumes were monitored by digital calipers twice a week and calculated using the formula: volume= (width)^2^ × length/2. At the experiment endpoint, mice were euthanized and tumors were harvested for *ex vivo* analysis and subsequent histology. All operations were approved by the Animal Ethics Committee of NCCS.

### Immunohistochemistry staining

Tumors were fixed in 10% buffered formalin overnight and embedded according to standard procedures. Tumor sections were cut to 4um thickness, mounted on glass slides, and air-dried at room temperature. After antigen retrieval, tumor sections were stained with antibodies and then with hematoxylin. Images of tumor sections were taken with a bright light microscope at X10.

### Statistical analysis

All statistical analysis in this study was performed using GraphPad InStat (GraphPad Software, CA, USA). Statistic significance was determined by two-tailed Student’s *t*-test in animal studies and error bars represent s.d. to show variance between samples in each group, or by one-sample t-test in other experiments and error bars represent s.d. to show variance between independent experiments. Supplementary information is available at Oncogene’s website (www.nature.com/onc).

## Author Contributions

J.Y. and J.H. designed the study; J.Y. and J.H. searched databases/literatures for kinase mutations in cancer genomes; M.B. prepared experimental materials; J.Y., W.H.N, Y.W, H.X., J.J.Y., and J.H. carried out molecular biology, biochemistry, and cell biology experiments; J.Y., P.L., and S.P.G. constructed mouse xenografts; J.Y., P.L., and A.L. performed immunohistology analysis; P.L., M.W., and J.H. supervised all experiments and interpreted experimental data; J.H. wrote the manuscript; M.B. and M.W. revised manuscript; and all authors commented and approved the manuscript.

## Acknowledgement

We thank the laboratories of Dr. Hui, Dr. Sabapathy, Dr. Virshup, and Dr. Anand for their help in experimental technologies. We also thank Dr. Andrey Shaw and Dr. Susan Taylor for their assistances. This study is supported by NCCRF startup grant (NCCRF-SUG-JH), NCCRF bridging grant (NCCRF-YR2016-JUL-BG1), NMRC seeding grants (NCCSPG-YR2015-JUL-14 and NCCSPG-YR2016-JAN-17), Duke-NUS Khoo Bridge Funding Award (Duke-NUS-KBrFA/2017/0003), Asia Fund Cancer Research (AFCR2017/2019-JH) and SHF Research Grant (SHF/FG692S/2016). The authors declare no competing interests.

